# Performance evaluation of inverse methods for identification and characterization of oscillatory brain sources: Ground truth validation & empirical evidences

**DOI:** 10.1101/395780

**Authors:** Tamesh Halder, Siddharth Talwar, Amit Kumar Jaiswal, Arpan Banerjee

**Author notes:** Authors have equal contribution (Tamesh Halder, Siddharth Talwar). Email address (Arpan Banerjee).

## Abstract

Oscillatory brain electromagnetic activity is an established tool to study neurophysiological mechanisms of human behavior using electro-encephalogram (EEG) and magneto-encephalogram (MEG) techniques. Often, to extract source level information in the cortex, researchers have to rely on inverse techniques that generate probabilistic estimation of the cortical activation underlying EEG/ MEG data from sensors located outside the body. State of the art source localization methods using current density estimates such as exact low resolution electromagnetic tomography (eLORETA) and minimum norm estimates (MNE) as well as beamformers such as Dynamic Imaging of Coherent Sources (DICS) and Linearly Constrained Minimum Variance (LCMV) have over the years been established as the prominent techniques of choice. However, these algorithms produce a distributed map of brain activity underlying sustained and transient responses during neuroimaging studies of behavior. Furthermore, the volume conduction effects, phase lags between sources and noise of the environment play a considerable role in adding uncertainty to source localization. There are very few comparative analyses that evaluates the “ground truth detection” capabilities of these methods and evaluates their efficacies based on sources in temporal cortex relevant for auditory processing as well as mesial temporal lobe epilepsies. In this Methods article, we compare the aforementioned techniques to estimate sources of spectral event generators in the cortex using a two-pronged approach. First, we simulated EEG data with point dipole (single and two-point), as well as, distributed dipole modelling techniques to validate the accuracy and sensitivity of each one of these methods of source localization. The abilities of the techniques were tested by comparing the localization error, focal width, false positive ratios while detecting already known location of neural activity generators under varying signal to noise ratios and depths of sources from cortical surface. Secondly, we performed source localization on empricial EEG data collected from human participants while they listened to rhythmic tone stimuli binaurally. Importantly, we found a less-distributed activation map is generated by LCMV and DICS when compared to eLORETA. However, control of false positives is much superior in eLORETA especially while using realistic distributed dipole scenarios. We also highlight the strengths and drawbacks of eLORETA, LCMV and DICS following a comprehensive analysis of simulated and empirical EEG data.

## 1. Introduction

Cortical oscillations play an important role in governing basic cognitive functions (Edelman and Mountcastle, 1978; Bressler and Kelso, 2001; Buzsáki and Draguhn, 2004). Several researchers have suggested that electromagnetic brain activity at specific frequency bands carries meaningful information about neural function, e.g. alpha waves at 10 Hz (Bollimunta *et al*., 2008; Llinás *et al*., 1999), beta at 15-30 Hz (Brovelli *et al*., 2004), gamma at 30 Hz and above (Bressler *et al*., 1993; Varela *et al*., 2001; Cheyne and Ferrari, 2013). Concurrently, time-locked transient responses have been useful for decades in electrophysiological research, both for understanding basic neurobiological functions as well as in clinical and other applications (Picton *et al*., 1974; Kutas *et al*., 1977; Pantev *et al*., 1995; Clark *et al*., 1995; Cheyne *et al*., 2006). Hence, identifying the neural generators of sustained cortical oscillations and task-specific transient neural responses from electro-encephalography/ magneto-encephalography (EEG/ MEG) is an extensive topic of research. Once identified with adequate reliability, the focal localization of sources will eventually reveal the underlying large-scale network governing cognitive tasks.

There are several source localization methods that exist in the literature, commonly known under the umbrella of inverse methods (Hämäläinen and Sarvas, 1989). Most of these techniques are based on fitting single/ multiple dipolar cortical source/sources within a defined cortical volume based on some assumptions about relationships between the sources (Hämäläinen and Ilmoniemi, 1994; Van Veen *et al*., 1997; Ishii *et al*., 1999; Gross *et al*., 2001; Hillebrand and Barnes, 2003; Liu *et al*., 2002; Sato *et al*., 2004). Some methods consider sources to have minimum correlation, e.g. synthetic aperture magnetometry (SAM) (Hillebrand and Barnes, 2003), linearly constrained minimum variance spatial filtering (LCMV) (Van Veen *et al*., 1997; Murzin *et al*., 2011). There are specialized measures that detect generators of oscillatory brain signals by considering maximum coherence between prospective sources, e.g., dynamic imaging of coherent sources (DICS) (Gross *et al*., 2001) and entropy based metrics (Lina *et al*., 2014). DICS is a frequency domain extension of beamforming methods over the initially developed time-domain beamformers, e.g., LCMV and synthetic aperture magnetometry (SAM) (Ishii *et al*., 1999) that are primarily used for determining the sources underlying time-locked ERP/ ERF components. In EEG, where deeper sources can affect scalp potentials, current density techniques such as minimum norm estimates (MNE) (Hämäläinen and Ilmoniemi, 1994) and exact low-resolution brain electromagnetic tomography (eLORETA) (Pascual-Marqui, 2007) has been the method of choice. Though, dynamic statistical parametric mapping (dSPM) (Liu *et al*., 2002) and sparse Bayesian learning (SBL) (Ramìrez *et al*., 2010) has been developed to improve upon the estimates of spatial filter detection, eLORETA is still by far one of the most robust methods for EEG source localization. eLORETA directly estimates current source density, a biophysically relevant parameter over a grid of plausible cortical locations for both detection of time-locked activity e.g., in event related potentials / fields (ERP/ERF) or frequency-locked activity e.g., spontaneous frequency bursts or steady state oscillatory responses to a periodic stimuli. Nonetheless, the source estimated by all methods are broadly influenced by depth, signal to noise strength of the neural activity, as well as the the correlation in the covariance of the signals (Belardinelli *et al*., 2012) and redundant informational content of high temporal resolution data. Often these manifest in distributed source activity estimation with diminished statistical power.

The accuracy of the location of neural activity along with lower false positives should be the expectation from any source localization technique. Many inverse methods can localize a transient or steady state response or both. However, the biological relevance or interpretation of these different information processing events can be very distinct. There are comparison studies that evaluate the performance across different methods e.g., Bradley *et al.* (2016); Hedrich *et al.* (2017); Jon Mohamadi *et al.* (2012); Mahjoory *et al.* (2017) or sometimes the performance of detecting a focal cortical source across modalities EEG and MEG Srinivasan *et al.* (2006); Mideksa *et al.* (2015). Few of them have directly compared current density measures and beamforming directly e.g. in Hincapie *et al.* (2017) and also very few work on ground truth validation (Herdman *et al*., 2003; Jon Mohamadi *et al*., 2011). In this article, we evaluate the performance of the key techniques eLORETA, DICS, LCMV and MNE on simulated EEG data. Since LCMV and DICS belong to the same class of beamformers, they were compared against eLORETA, to quantify localization efficiency between beamformers vs current density measures. We compared the results from eLORETA and DICS on a paradigm of evoked 40-Hz auditory steady state responses (ASSR) and eLORETA, MNE and LCMV for detecting the source of N100 activity when the same data was epoched time-locked to the stimulus onset. Our focus was to compare the specificity and sensitivity of some of the prominent algorithms primarily chosen based on their conecptual difference current density estimate vs beamforming to provide a basis for choosing one above the other when faced with the issues of transient or steady state response. Very rigorous comparison metrics e.g. localization error, spatial spread and false positive percentage were used to evaluate accuracy and sensitivity of results along with a evaluation of the performance of these methods at different depths of dipole placement in the simulated EEG data. The rest of this paper is organized as follows. In Section II, we introduce the simulation framework for testing the four methods (i.e., eLORETA, LCMV, DICS, MNE) on synthetic EEG data. We also outline the experimental design used to generate empirical EEG data on which we wish to compare eLORETA, LCMV and DICS. In section III we present the comparative results of applying eLORETA, LCMV, DICS and MNE on simulated and empirical EEG data. Finally in section IV we outline the comparative benefits and pitfalls of these methods.

## 2. Materials and Methods

### 2.1. Generation of synthetic EEG data

To localize the oscillatory activity, as well as the transient response, we simulated a time-varying sinusoid at 40 Hz and a mixture of Gaussian pulses, respectively. Both generated signals were free of noise, and were consequently added to the acquired empirical baseline, for realistic noise simulation. The magnitude of the dipolar source dynamics in cortical locations are represented by

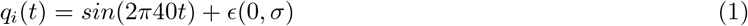

where *q_i_*(*t*) is the electric dipole moment at location *i* and at time *t, ϵ* is white noise with zero mean and standard deviation *σ*. We compared 3 conditions with respect to number of sources, by placing single point dipole, two-points dipoles and distributed dipoles in a MNI brain template according to the Montreal Neurological Institute (Mäller and Weisz, 2012). Single point source was placed at around the superior temporal region, in the left hemisphere (MNI coordinates: (−60, −28, 6)). Two sources were placed, one in the left hemisphere (MNI coordinates: (−60, −28, 6)) and the other in the right hemisphere (MNI coordinates: (64, −24, 6)) around the superior temporal region. Approximately, hundred point sources were placed within a spherical volume with radius of 12 mm in the left hemisphere centred around superior temporal region at (−60, −28, 6), according to brain template. Another set of hundred point sources were placed around right hemisphere auditory cortex seed area at (64, −24, 6), defining the distributed source condition. The resolution of the grid chosen for dipole simulation was 5mm and “*ft_prepare_leadfield.m*” code of FieldTrip toolbox was used for this purpose. Dipole moment orientations were assumed to be along the radial direction with respect to the BEM surface, to retain simplicity. We computed the scalp potentials for EEG at realistic sensor locations by applying a forward model (Mosher *et al*., 1999; Baillet *et al*., 2001) with realistic headshape using “*ft_dipolesimulation.m*” of the FieldTrip toolbox.

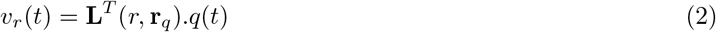

where, *υ* is the electric potential at sensor location *r, r_q_* represents all source locations, **L** represents the “lead field kernel”, (.)^*T*^ represents transpose and *q(t)* is the dipole moment. Synthetic EEG data were generated by varying signal to noise ratio (SNR) at the source space. Physiological SNR was estimated using a statistical measure, 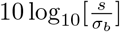, where *s* is peak-to-peak amplitude of EEG data during rhythmic auditory stimulation (see Experimental Methods below) and *σ_b_* is the standard deviation of the baseline data. We chose a wide range of SNRs (19 22 25 28 31 dB) to simulate mixture of Gaussian pulses mimicking transient reponse, both above and below the estimated physiological SNR level (25 dB), to allow us to evaluate the sensitivity of each method. Further, we introduced time lags between the signals generated from the left and right hemispheres for two and distributed dipole models. Time delays of: 0, 15, 30, 45 msec were added to the Gaussian pulse generated from the right hemipshere. Fig 1 shows simulated EEG activity on scalp surface with bilateral auditory cortical sources. Following Goldenholz *et al.* (2009), SNR was computed in decibel(dB) using the following equation.

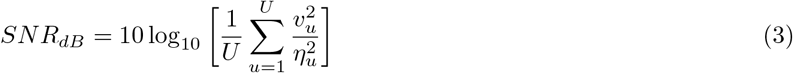

where *U* is total sensor count, *υ* is the signal on sensor *u* ∈ (1, 2, ⋯ *U*) provided by the forward model for a source with unit amplitude. The sensor space variance 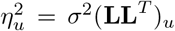. Therefore, for each time lag scenario, we simulated the Gaussian signal of 5 SNR values. Additionally, we also simulated sinusoidal signals mimicking oscillatory activity with different values of power spectra at 40 Hz. The power values were chosen with respect to the power spectrum computed for the empirical binaural condition data, such that the power of simulated sinusoidal at 40 Hz was 50%, 75%, 100%, 125%, 150% of the power at 40 Hz of the binaural condition, illustrated in Fig 1. Further, phase lags were introduced between the signals generated in left and right hemisphere: 0, *π*/2, *π* and 3*π*/2. Therefore, all the power ratio scenarios were computed for each phase lag condition.

**Figure 1:**
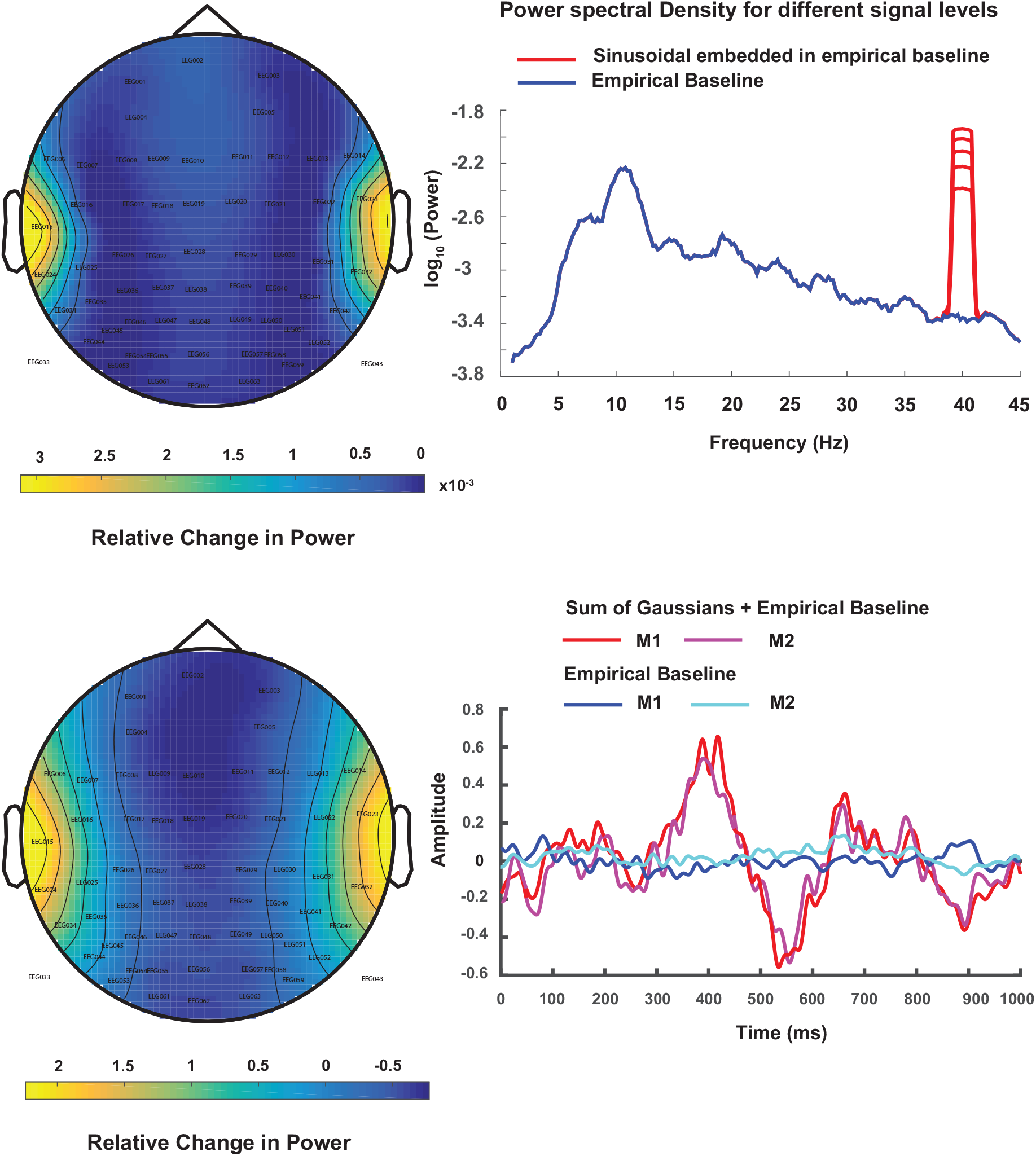
a) Topoplot of the difference in peak of spectral power at 40 Hz obtained from simulated EEG data where dipolar source time series was represented by a sinusoidal signal with frequency 40Hz embedded in empirical resting state EEG and empirical resting state EEG as baseline. Spatially averaged power spectra obtained from averaging the channel-by-channel spectrum from hypothetical scalp sensors are plotted in logarithmic scale. The time series on the scalp were obtained by applying forward modeling techniques on dipolar sources at auditory cortex locations using the boundary element method (BEM) b) Topoplot of the peak of difference signal when a mixture of Gaussian pulses was used to simulate ERP and empirical resting state EEG as baseline in BEM model as described in (a). The time series for dipole dynamics are plotted at hypothetical M1 and M2 sensors located near to the auditory cortices. The positive peak at 400 ms was used for generating the topoplot.

### 2.2 Source localization methods

The basic goal of any source localization technique is to compute the dipolar source locations and strengths inside the brain from measurements on the scalp (inverse of equation 2). In other words the objective is to estimate the spatial filter **W**_*S*_ from the relation

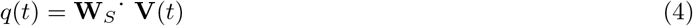

where, *q*(*t*) is the dipole moment at time *t*, **W**_*S*_ is the spatial filter matrix, and **V** is vector representation of all sensor time series. Obviously, the system of equations represented by equation 4 is ill-posed as number of sensors (dimension of vector **V**) are finite, but number of dipoles are unknown. So, different source localization methods attempt to estimate the **W**_*S*_ using diverse constraints posed by anatomy of the brain and functional relationships among brain areas during ongoing task.

#### 2.2.1 LCMV

Linearly constrained minimum variance (LCMV) belongs to the class of “beamformer” methods that enhances a desired signal while suppressing noise and interference at the output array of sensors (Barnes and Hillebrand, 2003). LCMV is built upon an adaptive spatial filter whose weights are calculated using covariance matrix of EEG/MEG time series data. A spatial filter computes the variance of the total source power which is allowed to vary but the output of the filtered lead field is kept constant. As a result the beamformer output is maximized for the target source but other source contributions are suppressed. LCMV attempts to minimize the beamformer output power

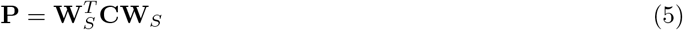

where **C** is the data covariance matrix. The entries to spatial filter matrix can be expressed as

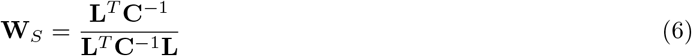

where *L* is lead field matrix, and following constraint is maintained **W**_*S*_ ̇**L**^*T*^ = 1.

#### 2.2.2 DICS

Dynamic imaging of cortical sources (DICS) beamformer (Gross *et al*., 2001) works with same constraint assumption of LCMV but extends the computation of spatial filter to the frequency domain. Here, sensor level cross spectral density (CSD) matrix replaces the covariance matrix and the spatial filter is applied to sensor level CSD to reconstruct the source level CSD of all combination of pairwise voxels. Hence, DICS directly estimates the interaction between sources at respective frequencies. The weight function can be written as

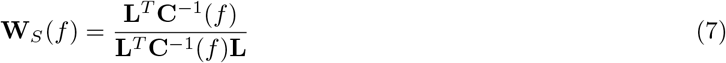

#### 2.2.3 eLORETA

Exact low resolution electrical tomography (eLORETA) (Pascual-Marqui, 2007) combines the lead-field normalization with the 3D Laplacian operator under the constraint of smoothly distributed sources. Compared, to DICS and LCMV where the constraint equation **W**_*S*_ ̇**L**^*T*^ = 1 is used, eLORETA seeks to minimize the product **H** = **W**_*S*_ ̇**L**^*T*^.

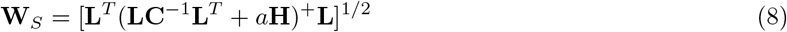

where ‘*a*’ is regularization parameter, + is the Moore-Penrose pseudo-inverse which is equal to the common inverse if the matrix is non-singular. **H** is also called the centering matrix or the surface Laplacian.

Low resolution imaging results in weak performance for recovering of multiple sources when the point-spread functions of sources overlap. Other methods have also tried to combine surface Laplacian with LCMV (Murzin *et al*., 2013), to estimate source-level connectivity.

### 2.3. MNE

Minimum norm estimates (MNE) has been a popular choice to localize evoked activity and tracking the distribution of the activations over a period of time. MNE is a distributed inverse solution that discretizes the source space into locations on the cortical surface or in the brain volume using a large number of equivalent current dipoles. It estimates the amplitude of all modeled source locations simultaneously and recovers a source distribution with minimum overall energy that produces observed sensor data consistent with the measurement (Hämäläinen and Ilmoniemi, 1994; Ou *et al*., 2009). The current density *q* can be calculated as

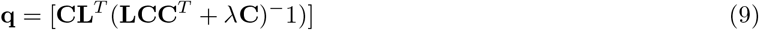

### 2.4. Measurements used for face validity of inverse algorithms

Using simulated data to test a method provides mechanism for ground truth validation. The exact location of a putative dipolar source is elusive in nature for real data, however, one can certainly set-up simulations when performance of a particular method needs evaluation. We employed three complementary measures to provide face-validity of the eLORETA in comparison with LCMV and DICS.

#### 1. Localization error estimation

Inverse methods estimate a cluster of point dipolar sources. To measure how much error is involved in source localization, we first computed the z-scores for all voxels. Further, we thresholded the -scores at 99.99th percentile and identified the cluster closest to the dipole location. Then we measured the Euclidean distance between the voxel with the maximum z-score of the nearest cluster source points and the actual source/dipole coordinates, to give us the ‘Localization Error’. This was done for each hemsiphere, separately. Consequently, the net localization errors are computed by summing up across two hemispheres and compared for eLORETA, MNE, LCMV and DICS.

#### 2. Degree of Focal localization

The size of a cluster in terms of sum of distances of all points from the voxel with the maximum z-score, gives a measure of focal localization of sources. Post finding the z-scores of all voxels, we thresholded the scores at 99.99th percentile and identified the cluster of source points closest to the dipolar location. Further, we computed the total sum of distances of each voxel in the nearest cluster, from the voxel in the cluster with the maximum zscore, as a measure of the spatial distribution of the estimated source. This gave us a quantitative approach to evaluate the degree of focal source localization. For practical reasons, cluster width computation was done for each hemisphere separately Murzin *et al.* (2013).

#### 3. Performance evaluation at various depths

The localization of deep sources has been the main factor limiting detection of true sources from M/EEG data. This is important since deep cortical areas constitute 30% of the cortical sources (Hillebrand and Barnes, 2002). We studied the effects of depth by positioning the dipoles at different distances from the auditory cortical locations mentioned earlier. The depth was varied along the x-axis, from 0 to 20 mm in steps of 1 mm, towards the center of the brain, in both hemispheres. Further, we computed the localization error and the focal width of the significant voxels obtained by localizing the dipoles placed at each depth. This was executed using the distributed dipolar method only. All simulated signals were added to the empirical baseline to retain the physiological SNR for the Gaussian pulses and the physiological power spectrum for the sinusoidal. The signals simulated were phase-locked (sinusoidal) and no time lags were added (Gaussian)

#### 4. Performance evaluation with various correlation

Correlation in the data covariance is an important variable which can influence localizing capabilities of beamformers (?). Therefore, we simulated multiple signals using the distribted dipolar model, consequently adding with the empirical baseline, such that there are various phase lags between the signal simulated in the left hemisphere and the right hemisphere. 4 phase lags chosen for frequency domain analysis were 0, *π*/2, *π*, 3*π*/2. The power ratios were matched as per the acquired empirical power ratios. 4 time lags chosen for time domain analysis were: 0, 15 30 45 msec.

#### 5. False positive percentage

We compared the “sensitivity” and “specificity” of eLORETA, LCMV and DICS, using ROC analysis (Metz, 1978). Here, we calculated the probability of incorrectly detecting an activation, also called ‘false positive (FP)’. Ideal detection should suppress FP. After thresholding the z-scores, we identified the number of significant clusters in each hemisphere, visually. Further we ran k-means clustering over significant voxel locations in each hemisphere and identified the nearest cluster to the *true* dipole location. Defining the nearest significant cluster/s from the dipolar location/s as the true positive/s, we further defined the false positive percentage by computing the ratio of number of significant voxels not present in the true positive (or nearest cluster) and the total number of significant voxels. We also compare the performance of all methods under parametric variation of signal to noise ratio at the source level and different kinds of source configurations, e. g., single point dipole, 2-point dipoles and distributed dipoles.

All analyses codes with simulated EEG data can be downloaded from the following weblink **https://bitbucket.org/sid03/source-localization/downloads/?tab=downloads**

### 2.5. Empirical EEG recordings

#### Participants

10 healthy volunteers (8 males, 2 females) aged between 22 – 39 years (mean age 28 years) participated in the study after giving informed consent, following the guidelines approved by Institutional Human Ethics Board at National Brain Research Centre. All participants were self-declared normal individuals with no history of hearing impairments, had either correct or corrected-to-normal vision and no history of neurological disorders.

#### Stimuli

Volunteers had to remain stationary in a seated position within a sound-proof room and hear auditory stimuli through 10 Ohm insert earphones with disposable foam ear-tips, binaurally for 200 seconds while fixating at a visual cross. Additionally they had a baseline block where they fixated at the visual cross for 200 seconds without any sounds being played. Sounds were pure tones of 1000 Hz frequency and 25 millisecond time duration, with 5% rise and fall times and were repeated with a frequency of 40 Hz during an ‘On’ block of 1 second duration interspersed between 2 ‘Off’ blocks where no auditory stimuli were presented. Stimuli were made using in STIM2 stimulus presentation system with audio box P/N 1105 at 85 dB.

#### Data Collection and Pre-processing

EEG data was acquired in an acoustically shielded room with 64 channels NeuroScan (SynAmps2) system with 1 KHz sampling rate. Brain Products abrasive electrolyte gel (EASYCAP) was used to make contact with scalp surface and the impedance was maintained at values less than 5*k*Ω for all volunteers. Baseline EEG data was recorded for 200 seconds with eyes open, no tone, and a fixation cross on a monitor in front of the participants. Baseline and binaural stimuli were presented while participants were asked to maintain fixation on the cross all along to reduce eye movements.

Recorded raw data were re-referenced with average reference and were detrended to remove linear trends from the signal. Epochs of 5 second duration were constructed by concatenating ‘ON’ blocks of 1 second each, after removal of an initial 50 seconds of the 200 seconds long session. This was done to capture auditory steady state responses (ASSR). Data were band pass filtered with cutoff frequencies 5-48 Hz, to concentrate on sources underlying ASSR.

For an evoked waveform analysis, after average re-referencing, epochs of 1 second duration of ‘ON’ blocks were extracted from the raw data during stimulus condition, then filtered with cut-off frequencies 0.5-48Hz, detrended and averaged across trials to generate the evoked potential. Thresholds of — 100μV and 100μV were used to reject blink-corrupted trials, meaning if at any point within the epoch, the voltage exceeded the threshold values, the entire trial was deleted from the subsequent analysis.

Sensor locations were taken from the template given in the fieldtrip toolbox. Colin 27 structural T1 was used for co-registration with the sensor locations for accurate source localization. A forward model was computed using Boundary Element Method (BEM) from the respective T1-image. For localization using the algorithms, we considered 0 % regularization for all methods. The ratio of source power between stimulus and baseline condition was calculated in each voxel, using (*Power*(*Stimulus*) – *Power*(*Baseline*))./*Power* (*Baseline*) for the current density measures and (*Power*(*Stimulus*) – *Power*(*Baseline*)) for the beamformers. After computing the source intensities in each volunteer, the individual grids were interpolated to the T1 image. The averaged voxel intensities across all participants were evaluated using non-parametric statistics and z-scores were computed for each hemisphere. The top 0.05% voxels were identified as sources.

## 3. Results

### 3.1. Simulated EEG data

Simulated EEG data was computed by placing electric dipolar sources at auditory cortical locations, according to single dipole, two-point dipoles and distributed dipole configurations, using equation 1 and projecting the source activity at realistic sensor locations of a Neuroscan (Compumedics Inc, USA) EEG cap using a realistic head model (equation 2, Baillet *et al*., 2001). We considered two types of temporal profiles for source activity, a sinusoidal signal mimicking the band-specific frequency response observed in typical EEG signal such as auditory steady state response (ASSR) and a mixture of Gaussian pulses representing the time-locked event related potentials (ERP). Baseline data was acquired empirically on which both the aforementioned signals were added. Two prototype examples of simulated scalp activity during task and baseline are illustrated in Fig 1. To observe the effects of correlation in the cross-spectral density matrix on source localization, phase lags were introduced to the simulated sinusoidal signals generated from each hemishphere (2 point and distributed dipoles), as well as, time delay was added to the gaussian pulses to the signal from the right hemisphere in 2 point and distributed models.

We applied eLORETA and DICS to perform frequency-locked source analysis on the sinusoidal data. Keeping the empirical baseline intact, we scaled the sinusoidal signal such that, we obtained different ratios of power of the sinusoidal at 40Hz, comparable to the values we found in our empirical data. Fig 2a, illustrates combined results from eLORETA and DICS algorithms on a brain surface rendered by the MNI brain, at power ratio of 1 (realistic SNR), for the distributed dipole model.

**Figure 2:**
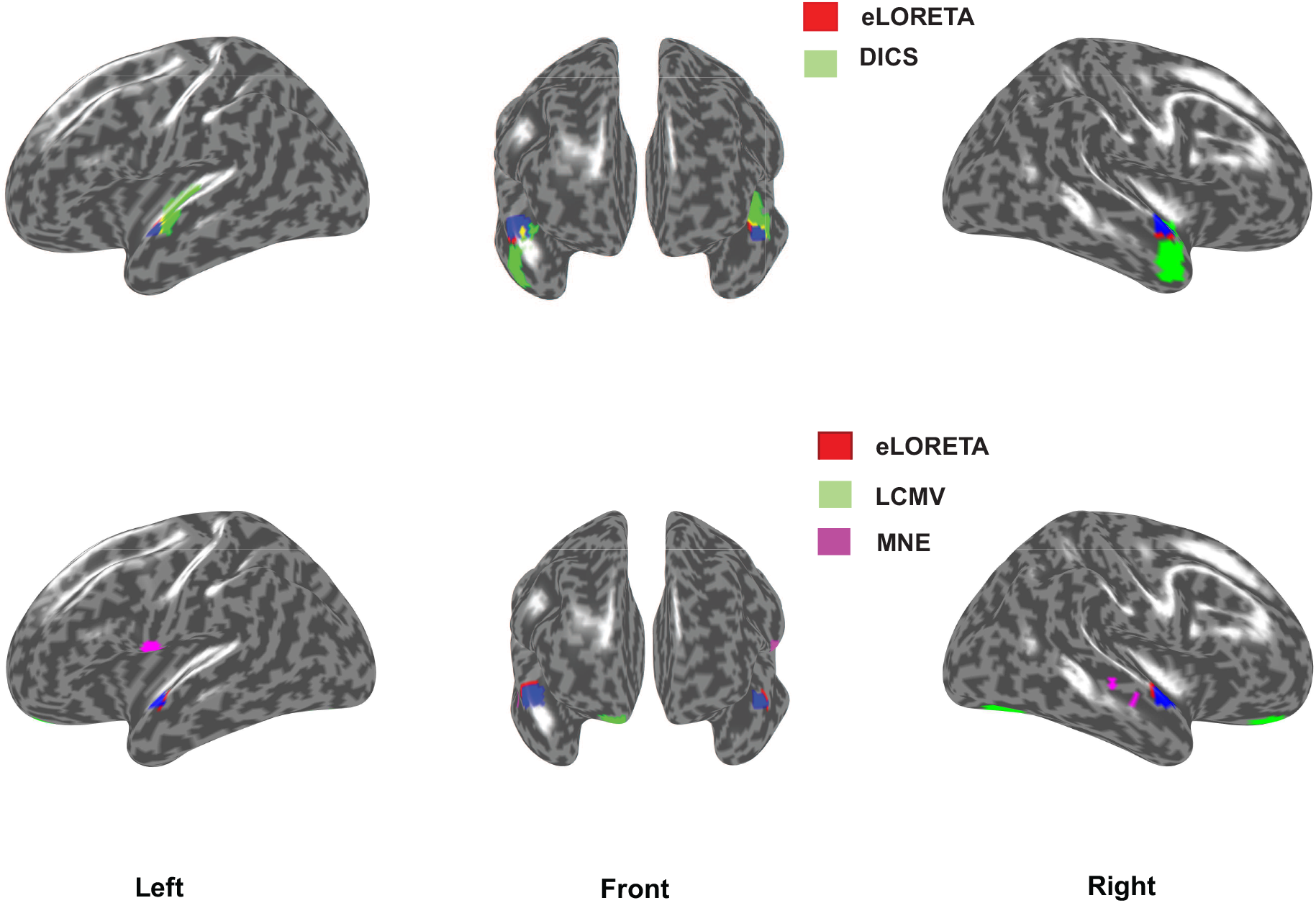
a) eLORETA source localization (red) vs DICS source localization (green) using frequency lock analyses on distributed dipolar source generated signals. eLORETA was applied to the simulated sinusoidal signal embedded in resting state EEG, where the simulated signals generated from each hemisphere had 0 phase lag. For DICS, the simulated signal from the right hemisphere had a phase lag of *π* with respect to the signal from the left hemisphere. The ratio of power spectrum at 40 Hz between the sinusoidal embedded in EEG and resting state EEG was chosen similar to the ratio found in our empirical results. b) eLORETA source localization (red) vs LCMV source localization (green) using time lock analysis. The results indicate 0 time lag scenario between the Gaussian pulses from each hemisphere. Physiologically realistic SNR 25 dB for simulated dipolar sources was chosen for this illustration.

The algorithms were employed for localizing the sources of the peak negative response in mixture of Gaussians signal, by selecting a time segment constituting of points ±25 ms around the peak (Fig 1b). For plotting activations, the source locations in the 3-D voxel space was projected to a surface plot using customized MATLAB codes (coord2surf.m in **https://bitbucket.org/sid03/source-localization/downloads/?tab=downloads**).

The localization error was computed by first defining each voxel as point in a cluster and thereby determining the distance of the voxel with the maximum z-score from the true dipole source in each hemisphere. For distributed dipolar sources, the centre of the spherically distributed source was considered as the true dipole location. The average of sum of distances from all such points to the voxel with the maximum z-score, normalized by the total number of points was used to quantify the focal localization of sources, in each hemisphere. All voxels were then transformed to their nearest projections on the cortical surface and identified as possible source locations in Fig 2. The quantitative evaluation of the performances of the inverse methods are addressed as follows.

#### Accuracy

Frequency analyses using eLORETA and DICS yielded similar localization errors with respect to different power ratios, in 1 dipole condition giving 0 false positives (Fig 3). However, eLORETA provided much lower localization error than DICS for 2 point and distributed dipole conditions. This was observed at lags: 0, *π*/2 and 3*π*/2, where DICS performed comparatively poorly. Interestingly, DICS performed better than eLORETA at phase lag of *π* in terms of accuracy, at which eLORETA’s accuracy deteriorated. Overall, the most significant observation was linked to the consistent performance of the algorithms regardless of the power spectra at 40 Hz.

**Figure 3:**
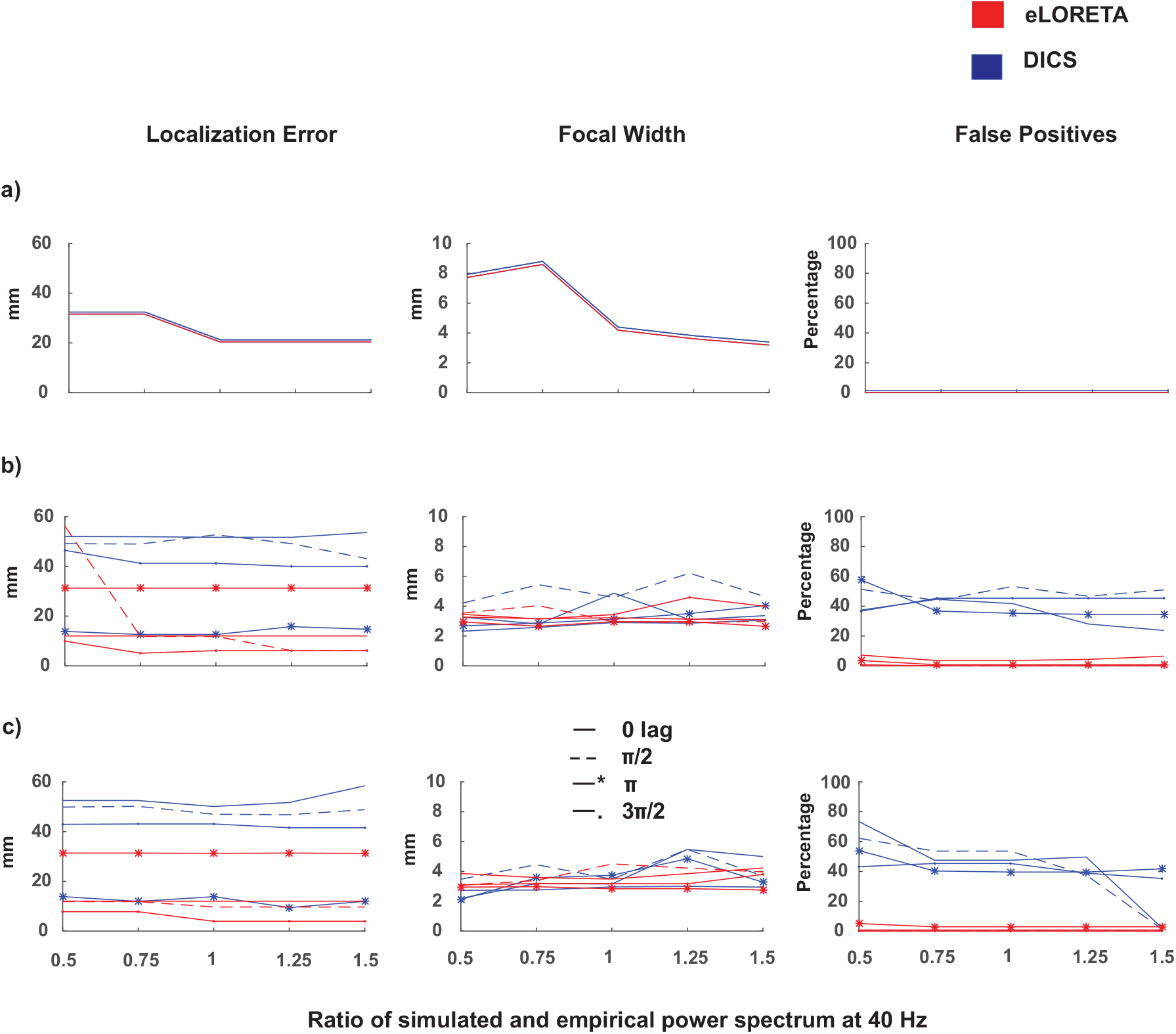
Localization error, Focal width and False positive percentage of (a) Single Dipole (b) Two Dipole and (c) Distributed dipole condition measured for eLORETA (red) and DICS (blue). Source localization was done for all power ratios (x axis) across different simulated phase lags of 0,π/2,π and 3π/2.

In time domain analysis, eLORETA performed better in all dipole configurations in contrast to MNE and LCMV (Fig 4). eLORETA consistently provided localization error of around 1cm or less for all time lags. LCMV’s accuracy with respect to SNR’s and time lags was not consistent as no trend could be observed. In contrast, MNE was observed to be better than LCMV at almost all scenarios with consistent errors across all SNRs. Therefore, it can be pointed that SNR and time lags affect the performance of the beamformers and minimal effects can be found in the current density measures, in terms of accuracy.

**Figure 4:**
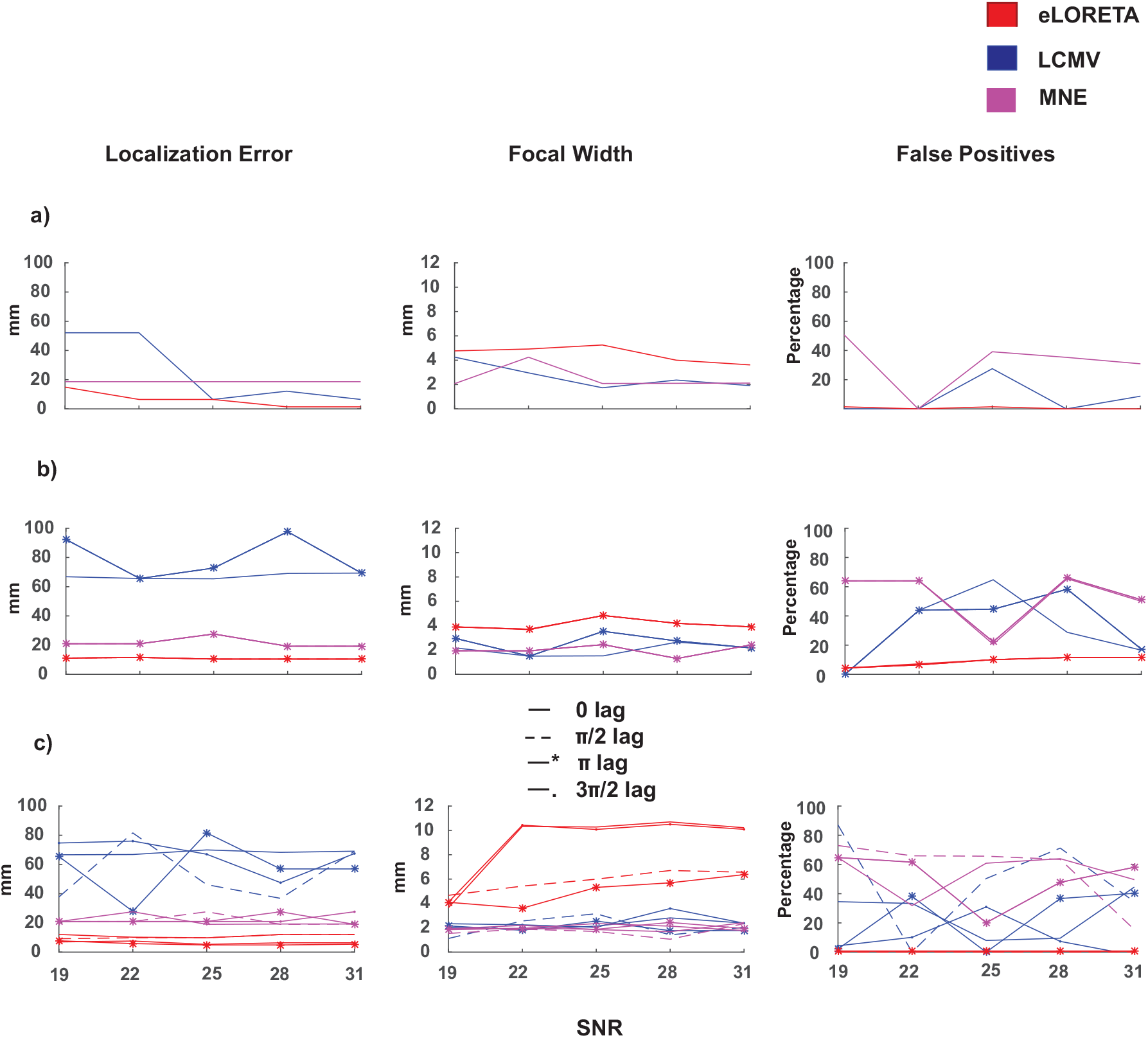
Localization error, Focal width and False positive percentage of (a) Single Dipole (b) Two Dipole and (c) Distributed dipole condition measured for eLORETA (red), LCMV (blue) and MNE (pink). Source localization was done for all SNR’s (x axis) across different simulated time lags where 25dB refers to the realistic scenario.

#### Localization Spread

eLORETA performance is observed to be comparable (similar focal localization) to DICS across all dipole conditions, as well as, power ratios (Fig 3). Apart from slightly higher values of focal localization for DICS at *π*/2 lag, both algorithms are efficiently focal.

The focal localization of eLORETA on timelocked signal was observed to reduce for single and 2 dipole condition, across different SNRs, however not varying across time lags (Fig 4). MNE and LCMV performance was comparable and also didnot vary across time lags. However, the focal width of eLORETA deteriorated in the distributed dipole condition and varied heavily across time lags, in contrast to unperturbed performance of LCMV and MNE. Additionally, the focal width of eLORETA increased with increasing SNR. However, it was noted that there was minimal effect of SNR on the focal width of the other algorithms.

#### Depth

To quantitatively measure the localizing capabilities of eLORETA, DICS, MNE and LCMV, source localization was executed for distributed dipoles at different depths. The localization error and the focal width were computed for the significant voxels, illustrated in Fig 5. The depth of dipolar locations was varied along the x-axis according to the MNI template, 1mm apart for each localization iteration. The deepest positions were selected as [−40, −28, 6] in the left hemisphere and [40, −24, 6] in the right hemisphere. The locations closest to the surface were chosen as [−60, −28, 6] in the left hemisphere and [60, 28, 8] in the right hemisphere.

**Figure 5:**
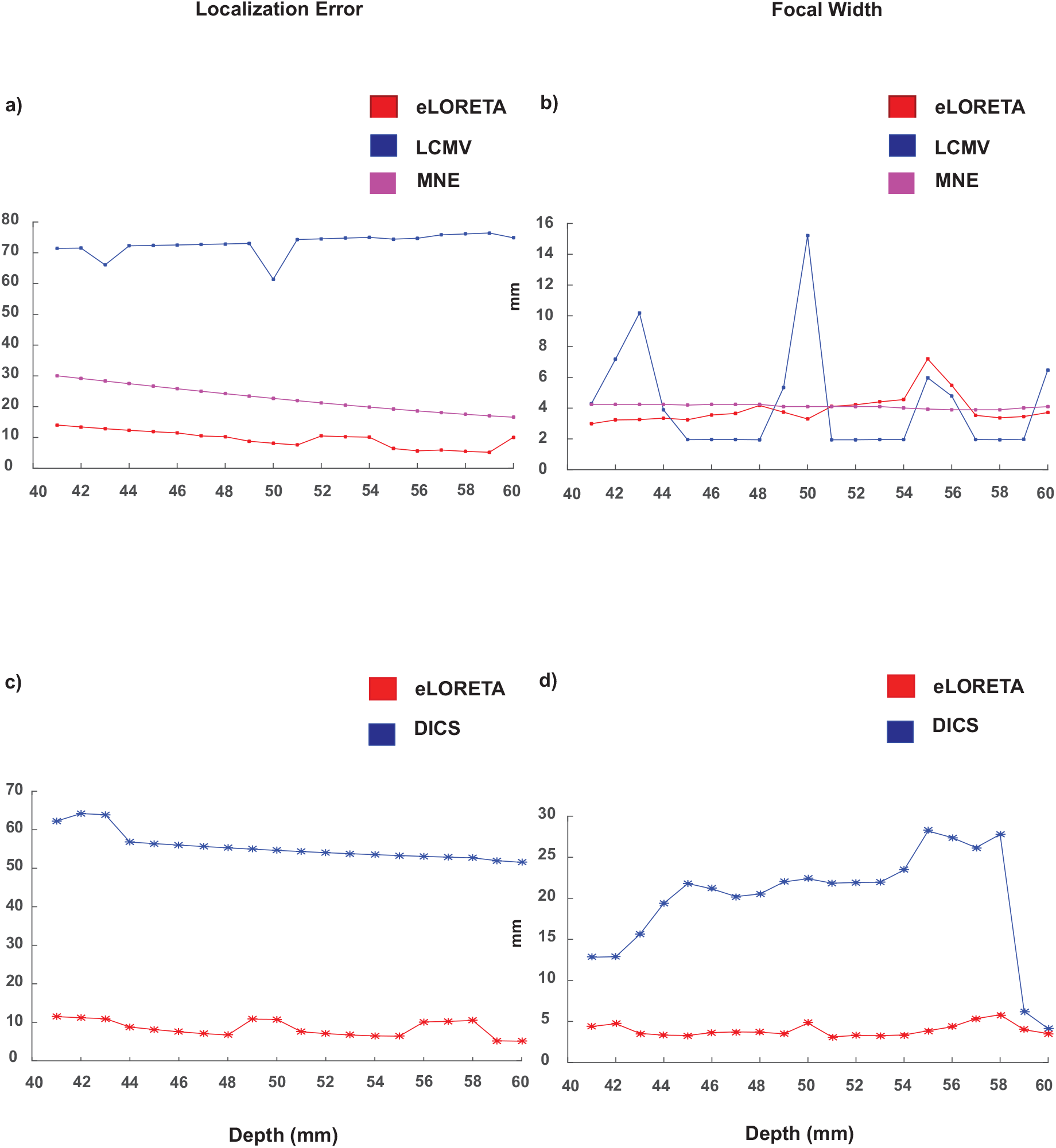
Localization errors and focal widths of all significant voxels computed for simulated distributed dipoles with the centre of the dipole cluster located at different depths from the auditory cortical locations. The depth of the dipoles decrease along the x-axis; the distance from the auditory cortex. (a) and (b) show the localization error and the focal width of the sources generated by the mixture of Gaussian signals respectively. (c) and (d) exhibit the localization error and focal width for frequency domain analysis.

The localization error for DICS declined and performed favourably as the depth of the dipoles decreases. A difference of 1 cm was observed between the deepest and the most superficial source for DICS. However, eLORETA gave consistent low localization errors across all depths. It is observed to be more focal than DICS across all depths, except for comparable focal width for superficial sources.

For the time-locked condition, the current density measures performed better than the beamformer, for localizing the Gaussian signal at all depths (performance of eLORETA>MNE>LCMV). Although LCMV was found to be more focal at certain depths, eLORETA and MNE were found to be focal across varying depths. eLORETA proved to be slightly more focal than MNE at deeper depths, in contrast to MNE being more focal the eLORETA at smaller depths. Therefore, eLORETA can be credited to have a higher degree of focal localization in comparison to MNE, LCMV and DICS, for localizing sinusoidal, as well as, Gaussian pulse, especially at deeper depths.

#### False positive percentage

To evaluate specificity, we computed the false positives after applying DICS, LCMV and eLORETA in different SNR scenarios, for all phase/time lags across different dipole conditions. All source points except the cluster of points nearest to the dipole location were considered as false positives (see section 2.4 for details).

With 0 false positives for 1 dipole condition, the false positives of DICS increased with increasing number of dipoles (Fig 3). With the percentage ranging from 30-70 percent, eLORETA provided a very high rate of true positives across all SNR’s and phase lags. Unsurprisingly, eLORETA gave a very low false positive fraction on localizing the Gaussian, similar to localizing the sinusoidal (Fig 4). MNE and LCMV gave high and comparable but consistent false positives fraction across all time lags in single and 2-dipole condition. However, in the distributed dipole case, the false positives of MNE and LCMV increased with varying time lags.

Table 1 classifies the performance of all the methods based on accuracy, localization and sensitivity across all dipole conditions for localizing 40 Hz sinusoidal signal (frequency analysis) and Gaussian pulse response (time-lock) respectively.

**Table 1:** Outcome of source localization performance based on different metrics for a) frequency analyses, eLORETA, DICS and b) time domain analyses, eLORETA, LCMV, MNE. If similar performance was achieved, both methods are mentioned with ‘/’.

| a) Frequency Analysis |  |  |  |
| --- | --- | --- | --- |
| Dipole Condition | Localization Error | Focal Width | False Positives |
| Single Dipole | eLORETA/DICS | eLORETA/DICS | eLORETA/DICS |
| Two-point Dipoles | eLORETA($0, \pi/2, 3\pi/2$)/<br>DICS($\pi$) | eLORETA | eLORETA |
| Distributed Dipoles | eLORETA($0, \pi/2, 3\pi/2$)/<br>DICS($\pi$) | eLORETA | eLORETA |
| b) Time-lock Analysis |  |  |  |
| Dipole Condition | Localization Error | Focal Width | False Positives |
| Single Dipole | eLORETA | MNE/LCMV | eLORETA |
| Two point Dipoles | eLORETA | MNE/LCMV | eLORETA |
| Distributed Dipoles | eLORETA | MNE/LCMV | eLORETA |
| — |  |  |  |

### 3.2. Empirical EEG data

#### Source localization underlying 40Hz EEG activity

The Fourier spectrum of each EEG channel time series were computed by multi-taper method with number of tapers = 2, using Fieldtrip function ft_freqanalysis.m. Power spectral density of empirical EEG data and the ERP time locked to the onset of a single tone stimulus are shown in Fig 6. In Fig 6a, the topoplot of the difference in power between binaural and baseline conditions at 40 Hz is shown along with the log of power across all trials. There are peaks at alpha band 8-12 Hz in both binaural and baseline conditions the difference between which was not significant (two-sample *t* = −0.018, *p* = 0.95) whereas the binaural condition had a sharp rise in power at 40 Hz which was significant (two-sample *t* = 0.18, *p* < 0.0001). The t-tests were performed on logarithm of power spectrums from two conditions within a frequency bands, 8 – 12 Hz for alpha and 39.8 – 40.2 Hz for evoked 40 Hz. Localization results are illustrated in Fig 7a. The left, top and right views of activation are shown where red regions represent sources from eLORETA analysis while green regions represent activation plots generated by DICS.

**Figure 6:**
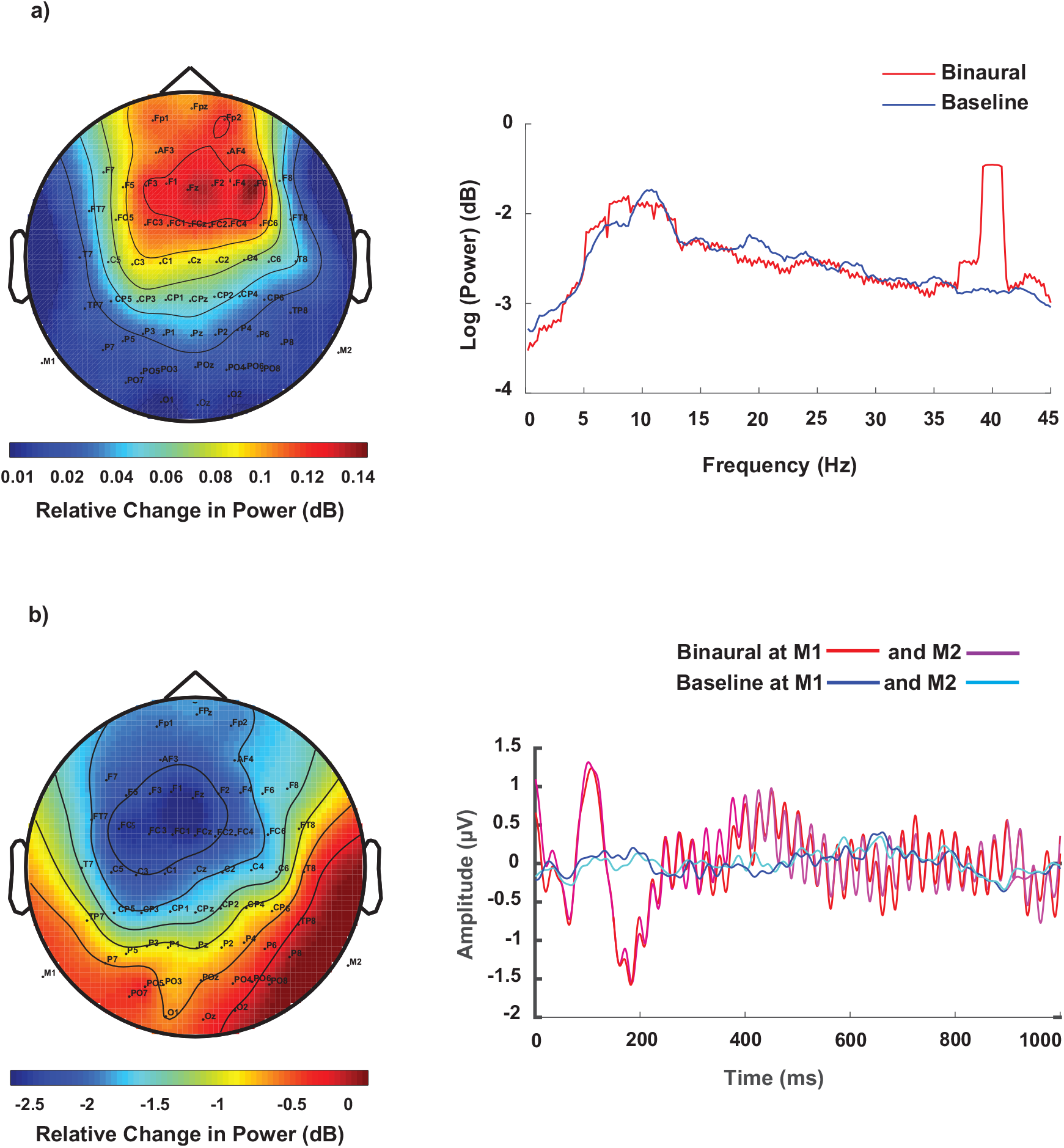
a)Topoplot of spectral power difference at 40 Hz and grand average of spectral power across all sensors, trials and participants in binaural and the baseline conditions. Power spectral density was calculated for 5 sec windows after rejecting an initial 50 sec out of total duration of 200 seconds for which the rhythmic tones were played. b) ERP responses of channels M1 and M2 across trials and participants for binaural and silent baseline conditions and the topoplot for the difference signal at the peak of N100 response (at 110 ms).

**Figure 7:**
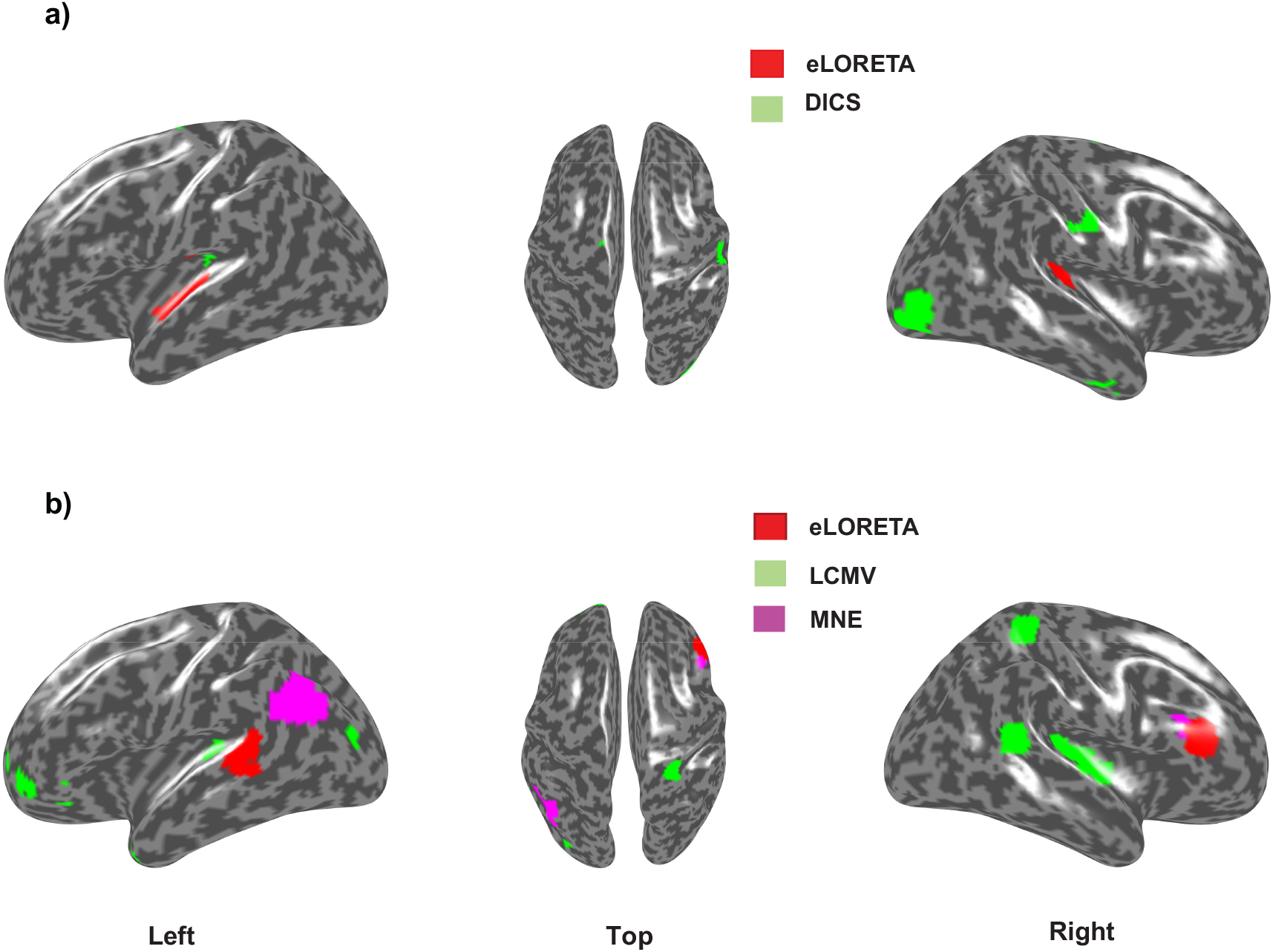
Left, top and right view of significantly active cortical sources underlying for a) 40 Hz auditory steady state response (ASSR) b) N100 component of the event related potential (ERP). The red colored regions show estimated sources from eLORETA while green regions show estimated sources from a) DICS and b) LCMV analysis. The pink regions in b) show MNE source localization results.

Acknowledging the absence of ground truth concerning true sources that exist in empirical data, we evaluated the focal width of all the significant clusters post thresholding the z-scores. 1 such cluster was found in each hemisphere around the auditory cortex for eLORETA, providing a mean focal width of 3.958*mm*. In contrast, DICS yielded 1 cluster at the auditory cortex in the left hemisphere and 7 distributed clusters in the right. The mean focal width across DICS clusters was 2.8324mm, lesser than that of eLORETA. Understanding that the thresholding can vary focal width and number of clusters, we maintained the same threshold for both the algorithms for a fair comparison. It is to be noted that the mean focal width of 2 clusters of eLORETA can be decreased with higher thresholding.

#### Source localization of N100 response

In Fig 6b we show the ERP responses to the binaural tone and the ERPs in baseline condition, averaged across all trials and participants (grand average). A negative peak around 100 ms post onset of tone stimulus (N100) was observed in the binaural condition with a latency of around 110 ms. The topoplot represents the spatial map of the difference in relative changes of amplitude between ERPs from the the binaural and baseline conditions across all channels and trials.

Next, we computed the underlying source activation during the N100 response using LCMV and eLORETA. In Fig 7b we plot the source activations (top 0.05% voxels similar to 40Hz case) in epochs of duration 50 ms, within which the 25th ms corresponds to the peak of N100. The beamformer localized bilateral auditory cortices, along with other distributed significant clusters. It can noted that LCMV may localize the underlying activity, however, has higher probability of yielding false positives, due to its distributed activations. In contrast, the current density measures localized the left auditory cortex (eLORETA) and left posterior superior temporal sulcus (MNE). Computing only 1 cluster for each hemisphere, eLORETA and MNE yielded a significant cluster in the right frontal regions. The focal width of LCMV (2.1706 mm) was lesser than current density measures (3.1-3.3 mm), similar to the case in the frequency domain analysis.

## 4. Discussion

Identifying the sources underlying key events of information processing such as ERP peaks or oscillatory brain activity such as spontaneous gamma oscillations are the objectives of many research studies. However, different inverse methods provide different solutions leading to no agreement of which algorithm is the ‘best method’, as we illustrated in this manuscript with simulated and empirical EEG data. Even though the selection of the best method can be guided by the nature of the hypothesis in a putative experimental design, a systematic comparative account of the efficacies of few prominent methods is currently missing in the literature. To address this issue we compared methods, eLORETA (Pascual-Marqui, 2007), LCMV (Van Veen *et al*., 1997), MNE (Hämäläinen and Ilmoniemi, 1994) and DICS (Gross *et al*., 2001) using the metrics that evaluates accuracy, sensitivity and specificity across three dipolar models (single, two-point and distributed) for the simulated sinusoidal signal (mimicking the steady state 40 Hz) and a mixture of Gaussian pulses (representing the time locked ERP). Furthermore, we chose an empirically observed baseline which is a key ingredient for every inverse method. The models were simplistic, however, since we knew the exact location of dipole/s, ground truth validation was possible. All methods are able to retrieve the location of the true dipolar sources for a physiologically relevant SNR and frequency power ratio (blue areas in Fig 2). Nonetheless, the study was conducted across different SNRs and frequency power spectra to test each method’s sensitivity and specificity to noise. Furthermore, performance of all methods were evaluated with respect to dipolar depth, phase lags among sources, accuracy and localization spread over space. Also, we conducted source localization by collecting empirical EEG data exhibiting 40 Hz ASSR (Fig 7). DICS and eLORETA were used to compute the sources underlying the 40 Hz activity and LCMV and eLORETA were used to compute the sources underlying the N100 response (Fig 7). Thus, we could outlay the hallmarks of each method in a organized framework.

A general consensus emerges from comparing the algorithms that there is no clear winner (Table 1) if accuracy as well as sensitivity and specificity are all taken together as guiding parameters. DICS gives better accuracy than eLORETA in single and two-point dipole conditions even at low SNRs, however, the focal width of eLORETA generated sources are always slightly better most likely due to the minimization of the surface Laplacian component while estimation of the spatial filter. For distributed dipole scenario, focal width of eLORETA results were very similar to DICS, in fact getting better with higher SNR. Interestingly, eLORETA shows significant control on the false positive ratio in the distributed dipole condition, proving to be the method of choice for estimating sources underlying frequency response in a more exploratory setting. This is indeed a very important point to note for increasing number of studies studying resting state functional connectivity (Canuet *et al*., 2011; Custo *et al*., 2017). Once a hypothesis is put in place with some prior knowledge about the involvement of prospective brain networks one can go for DICS that can produce more accurate results (Tan *et al*., 2016). Interestingly, DICS results in lowest localization error with maximum phase lag of *π*, whereas the effect was reverse for eLORETA.

For source localization of ERP peaks, eLORETA majorly yielded better accuracy and specificity (in terms of favorable false positives) than LCMV and MNE, even though the focal width of eLORETA sources were considerably larger than LCMV and MNE. The pitfalls and advantages of each method are summarized in Table 4. As in case, of frequency domain analysis, we would recommend eLORETA for a exploratory level analysis whereas the LCMV or MNE for more hypothesis driven identification of sources.

An alternative solution to increase the probability of isolating an active source is to combine 2 or more methods and take the overlap of sources detected from them as the plausible source configuration. We provide a blueprint of these using our empirical and simulated data. Both spectral domain eLORETA and DICS were able to pinpoint the left auditory cortex, the location of one true source (Fig 2). Even though both of the methods overlap at some spurious locations in frontal and temporal cortices, nonetheless the number of false positives obtained with two methods combined are drastically low. This gives us the confidence that the ASSR involves strong activity in primary sensory regions of auditory processing e.g., bilateral middle temporal gyrus as reported by earlier studies (Herdman *et al*., 2003; McFadden *et al*., 2014; Tan *et al*., 2016) and as we observe in Fig 7. Similar proof-of-concept illustration is available for time domain analysis as well where we combined eLORETA and LCMV to identify the sources underlying N100 peak. Obviously, validation from 2 or more methods will give a strong confidence in the result of source localization.

In future we think an overlap-approach might result in focal localization with minimum number of false positives. This will particularly benefit the identification of sources whose activity may be relevant for a particular context. For example cross-frequency coupling (CFC) between alpha-gamma rhythms are being postulated to be important for gating of attention (Klimesch, 2012). How to identify a cortical subnetwork whose nodes show CFC out of the whole alpha and gamma networks is an important methodological challenge. We believe a conjunction of methods strategy to identify the potential sources will be crucial from the perspective of reliability as well as accuracy. Thus our study provides a blue print for employing source-localization techniques to isolate more subtle features of signal processing.

## Acknowledgement

This study was funded by Ramalingaswami fellowship (BT/RLF/Re-entry/31/2011) and Innovative Young Bio-technologist Award (IYBA), (BT/07/IYBA/2013) from Department of Biotechnology, Ministry of Science & Technology, Government of India. AB acknowledges the support of NBRC Core funds for research and Distributed Informatics Centre, NBRC for infrastructural stupport. We also thank Dr Dipanjan Roy for helpful comments in an earlier version of the manuscript.

